# Including Phylogenetic Conservatism of Shortgrass Prairie Restoration Species Does Not Improve Species Germinability Prediction

**DOI:** 10.1101/2022.07.12.499320

**Authors:** Yanni Chen, Dylan W. Schwilk, Robert D. Cox, Matthew G. Johnson

**Affiliations:** Department of Biological Sciences, Texas Tech University, 2901 Main Street, Lubbock, Texas 79409, USA; Department of Natural Resources Management, Texas Tech University, Lubbock, TX 79410, USA

**Keywords:** ecological restoration, seed germinability prediction, phylogenetic comparative method, phylogenetic conservatism, phylogenetic signal

## Abstract

**PREMISE:** We investigated whether phylogenetic conservatism can improve the performance of seed germinability prediction models. Previous studies in tallgrass prairie and alpine meadow revealed that seed morphological traits demonstrate phylogenetic conservatism. We hypothesized that phylogenetic conservatism in seed traits could help predict the seed germinability, under the assumption that seed traits contain phylogenetic signals.

**METHODS:** We measured seed germination percentage and seed morphological traits (seed mass, seed height, and seed surface area) on 34 native species from shortgrass prairie in North America. We supplemented these data with similar data from the literature on 11 more species. We calculated the robustness of the phylogenetic signal of each trait to the number of species sampled. We also compressed the phylogenetic distance matrix to a two-dimensional space, and applied the Akaike information criterion to evaluate the effects of phylogeny on seed germinability prediction models.

**KEY RESULTS:** We found weak but significant phylogenetic signals in seed mass and seed height in the full data set. These phylogenetic signals were not able to improve seed germinability prediction model performance among shortgrass prairie species. Our robustness tests of phylogenetic signals using random sub-sampling showed that the detection rate of phylogenetic signals in seed mass was increased along with the expansion of species pool, and nearly 100% at 40 species. However, the detection rate of phylogenetic signals in seed height was constantly low, around 20%.

**CONCLUSIONS:** When the phylogenetic signals are weak, the phylogenetic position does not improve germinability prediction model performance. Therefore, phylogenetic signals detected during a single species pool calculation may not accurately reflect the phylogenetic conservatism of the trait in a plant community. We suggest testing for robustness of phylogenetic signals using random sub-sampling tests.

## Introduction

The need for ecological restoration is constantly increasing. For example, the September 2014 United Nations Climate Summit suggested the need for 350 million hectares to be restored worldwide by 2030 (Bonn Challenge, https://www.bonnchallenge.org/). Tremendous numbers of native species will be needed to meet this need. Most ecological restoration projects select only a small number of species out of the community species list to conduct ecological restoration (Kiehl *et al*. 2010). Given the low numbers of species selected for any specific restoration project, maximizing the benefit from selected species is key. Thus, ensuring that the selected species have high final germination percentages is a high priority because seed germination ranks as one of the top restoration challenges (Larson *et al*. 2015). Therefore, lab assessment formulas to narrow down the restoration species list could aid species selection in many restoration projects.

Seed dormancy regulates seed germination but is complicated and hard to predict. In over 90% of species, seeds dry and start primary dormancy by the time of harvest (Finch-Savage and Leubner-Metzger 2006; Subbiah *et al*. 2019). After dispersal, seeds can have secondary dormancy, a shallow physiological dormancy which is broken by responses to environmental cues (Finch-Savage and Leubner-Metzger 2006). Multiple categorical seed dormancy types are widely represented in plant species, including morphological dormancy (MD), physical dormancy (PY), physiological dormancy (PD), and morphophysiological dormancy (MPD; (Baskin and Baskin 1998). Physiological dormancy is thought to be the ancestral state of seed dormancy and also serves as the diversification hub for different dormancy types (Willis *et al*. 2014). Considering the complexity of dormancy stages and the lengthy experiments needed to distinguish these types (Finch-Savage and Leubner-Metzger 2006), it is desirable to predict seed germinability success through other related traits.

Low germination rate hinders restoration and, given limited resources, managers desire to only include species with predictably high germination rates. Several seed traits are related to seed germination and might serve as more easily measured predictors of final germination percentage. In general, mass is a good indicator of seed germination, as small seeds tend to germinate faster (Westoby *et al*. 2002a; Barak *et al*. 2018), while large seeds can stay dormant longer and produce stronger seedlings after germination (Leishman *et al*. 2000; Westoby *et al*. 2002a). The rationale behind this phenomenon is related to nutrition stored in the seed under either a “larger-seed-later-deployment” interpretation (Ganade and Westoby 1999; Leishman *et al*. 2000; Kidson and Westoby 2000) or “cotyledon functional morphology” hypothesis (Hladik and Miquel 1990; Kitajima 1996a, b). Furthermore, seed size and seed shape are also traits influencing seed germination by stimulating or delaying seed germination through wind, water, or animal dispersal (Howe and Smallwood 1982). Large seeds generally have advantages for dispersal related to entrapment strategies, such as net trapping, surface tension, and wake trapping (Jager *et al*. 2019), especially for wind-dispersed species (Zhu *et al*. 2019). Specifically, seed morphological traits influence both seed primary dispersal (seed departure from parent plants) and secondary wind dispersal (seed lifting off the ground by wind power) (Zhu *et al*. 2019). Primary dispersal is mainly driven by dispersal height and terminal falling velocity, which are influenced by seed morphology (Sheldon and Burrows 1973; Jongejans and Telenius 2001). Secondary dispersal distance strongly depends on the lift-off velocity, which is influenced by seed height and seed surface area (van Tooren 1988; Schurr *et al*. 2005; Zhu *et al*. 2022). There are many other seed physiological traits associated with seed germination that are not commonly tested, such as base water potential, cardinal temperature, thermal time and hydrothermal time for germination (Bradford 2002; Hardegree *et al*. 2013).

Seed germination trials are time consuming, therefore, predicting germinability for species without conducting such trials could benefit restoration. Seed morphology traits are potential predictors of germination rate. If dormancy and lack thereof are evolutionarily conserved, then it may be possible to predict seed germination rate of unmeasured species based on the rates of closely related taxa. A phylogenetic tree models the inferred evolutionary branching history of a group of taxa (Baum and Smith 2013). A phylogenetically conserved trait will tend to be most similar among species close together on the phylogenetic tree. The common test for such phylogenetic signals is Blomberg’s K (Blomberg *et al*. 2003, Revell 2008), but it is also possible to include all pairwise phylogenetic distances among taxa in linear models through the method of phylogenetic residuals (Revell 2010). Phylogenetic trait conservatism is common across many traits and clades (Bu *et al*. 2016, Barak *et al*. 2018, Duncan *et al*. 2019). Adding phylogenetic residuals to the generalized least square model can take the evolution of unmeasured traits into account and improve the prediction model’s accuracy. This work has two major goals. The first goal is to test whether adding phylogenetic information among species (presented by x-y coordinates transferred from phylogenetic tree topology) can improve predictions of germination rate based on seed morphology. Adding phylogenetic information might improve predictions if the germination rate shows a phylogenetic signal or if the seed morphology effect on germination rate interacts with phylogeny. There is precedent for using phylogeny for this purpose: In a study of species native to tallgrass prairie, Barak et al. (2018) confirmed that adding the phylogenetic residual improved the accuracy of the seed germinability prediction model due to the phylogenetic conservatism in both seed germination and morphological traits. However, because phylogenetic tools are unfamiliar and inaccessible to restoration practitioners and due to a historical separation between evolutionary biology and applied ecology, phylogenetic methods have not been broadly applied to restoration practice (Hipp *et al*. 2015).

The second major goal of this work is to determine how the size of a sample of taxa from an ecological community influences the power to detect phylogenetic signals in traits. The sample size and combination of given species influence tree topology and branch length during phylogenetic signal calculation. In empirical examples, the detection of phylogenetic signals is strongly related with the number of species included, with twenty or more species usually considered sufficient for estimation of Blomberg’s K (Blomberg *et al*. 2003). However, in phylogenetic comparative analysis aimed at answering evolutionary questions, the combination of species is commonly fixed. For applied restoration use, the practitioner will need to measure traits on some sample of species from a particular community. By examining how this sample of taxa influences the calculation of Blomberg’s K, we aim to provide guidelines for estimating the robustness of this calculation.

To address these two major goals and test the potential role of phylogeny for improving restoration practice, we asked four research questions: 1) Do seed traits and seed final germination percentages exhibit phylogenetic signals? 2) Among seed traits, which one is the best predictor of seed final germination percentage? 3) Does including phylogenetic residuals improve the seed germinability prediction? 4) Do the sampling size and species composition influence phylogenetic conservatism detection in shortgrass prairie species?

## Material and Methods

To determine the relationship between seed germinability and seed morphological traits, we measured seed germination percentage, seed mass, seed height, and seed surface area in 45 species which are native to the shortgrass prairie of North America (Table 1, Figure 1). All of our raw data and calculations were demonstrated in our interactive Shiny Application (Figure 2, https://chenyanniii.shinyapps.io/Phylo_Compar_Traits/).

**Table 1.**
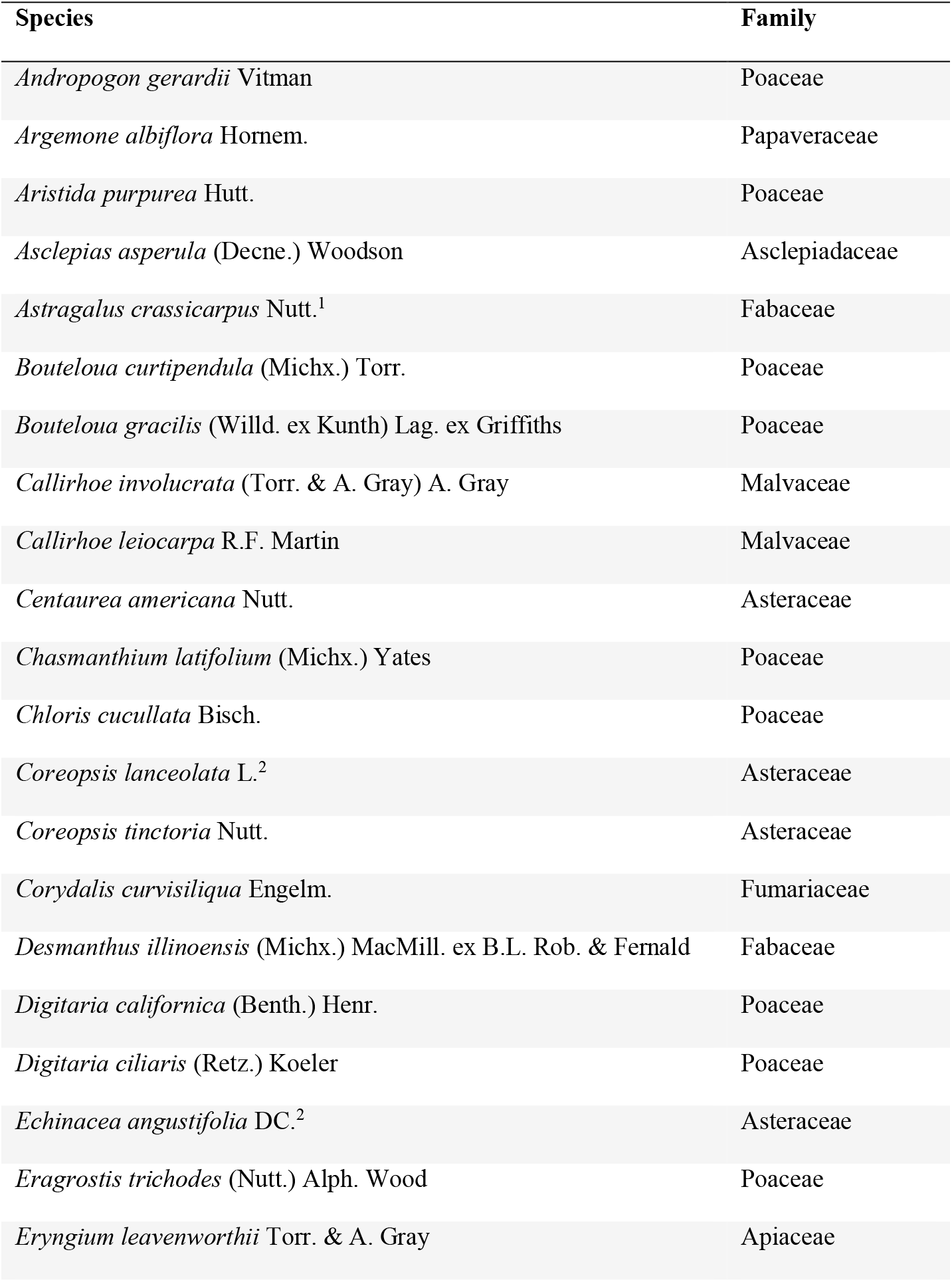
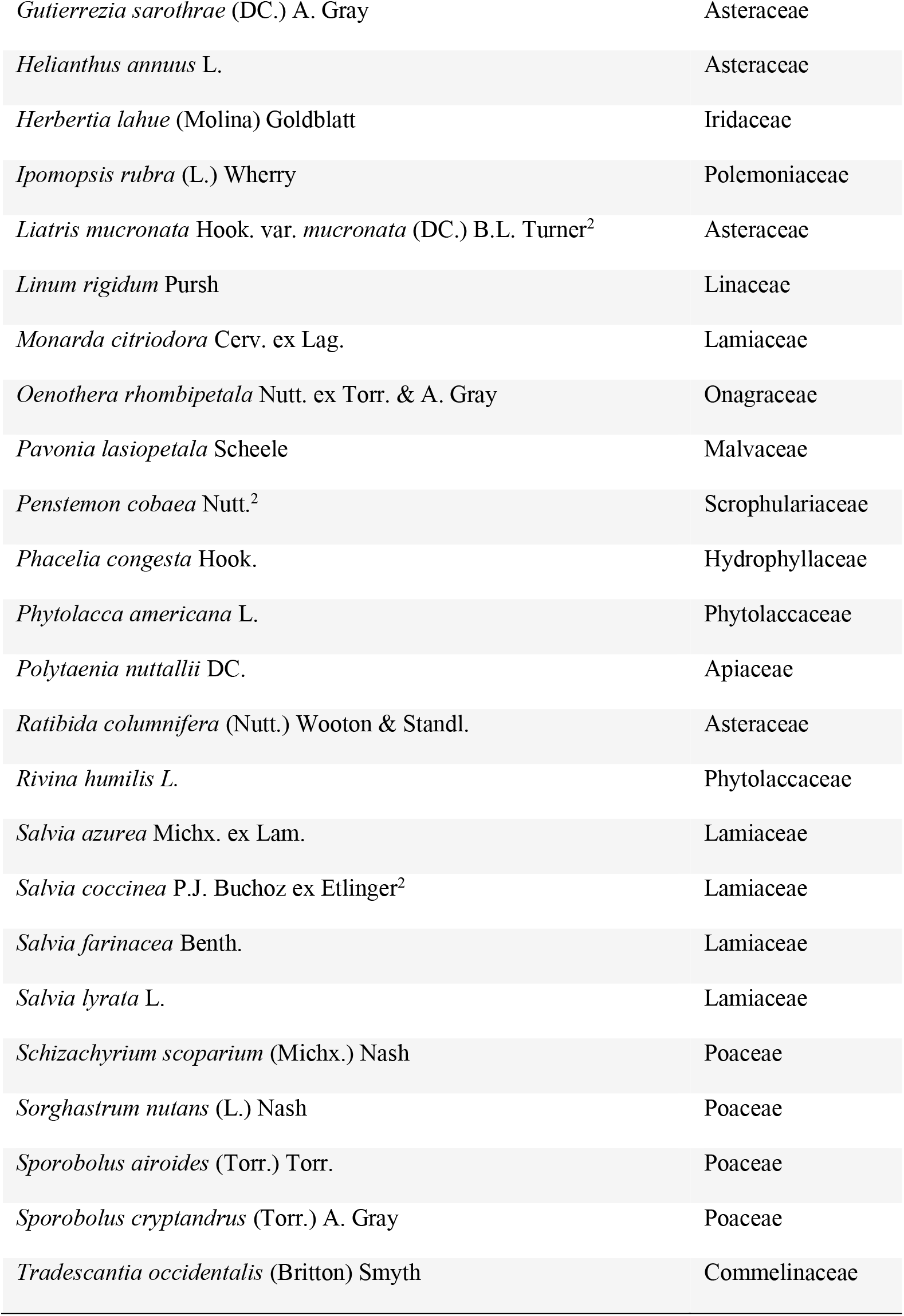
Forty-five native species were selected in this study, which are commonly involved in restoration practice and range management in shortgrass prairie. Most of the species were bought from Native American Seed, tested in controlled environments, 6 species were cited from (Chou *et al*. 2012)^1^ and 5 species were cited from (Schwilk and Zavala 2012)^2^.

**Figure 1.**
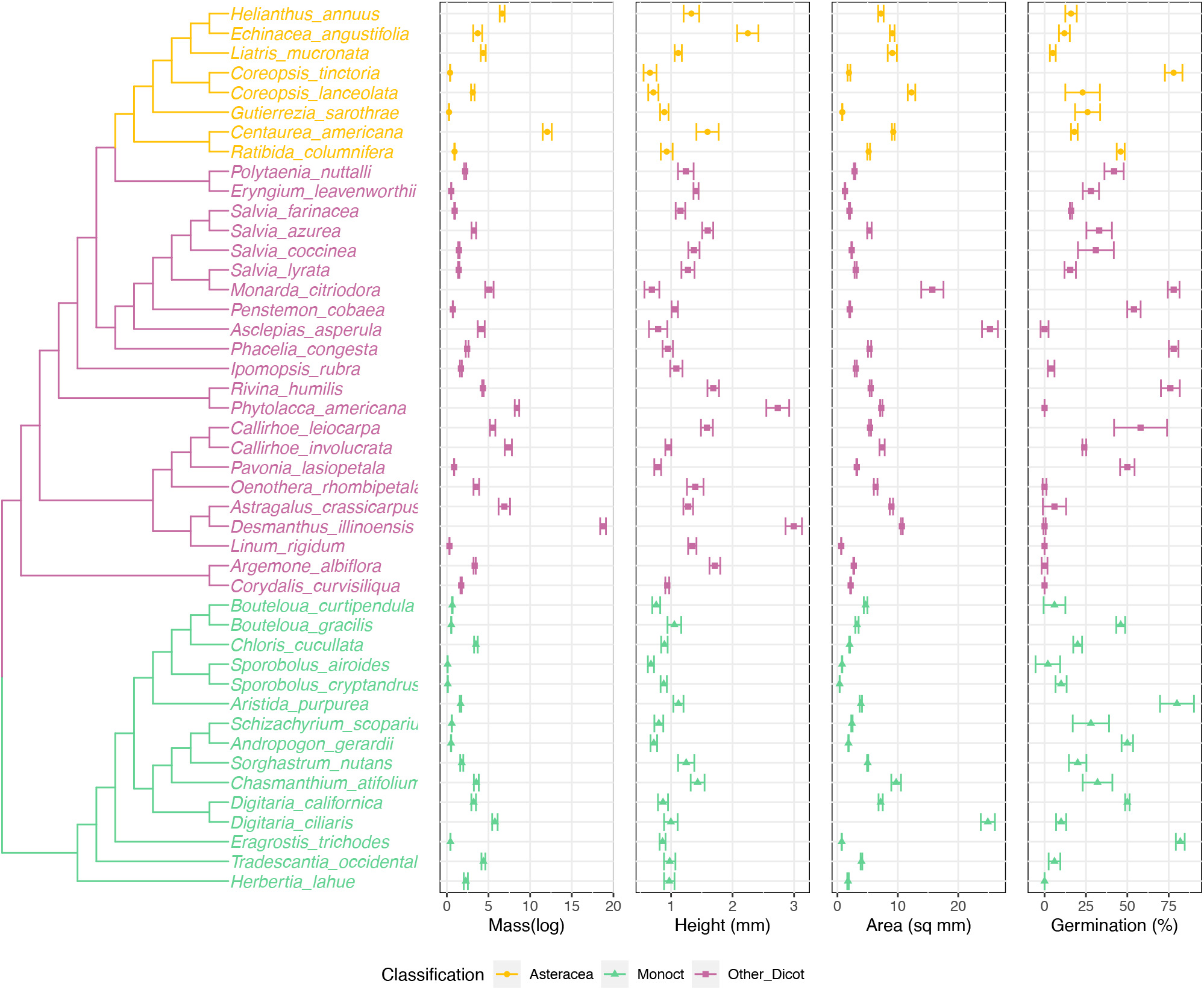
Phylogenetic tree of species and species seed traits values (seed mass, seed height) distribution along the phylogenetic tree. Phylogenetic tree was generated from the pruned Zanne et al. tree (Zanne *et al*. 2014), including 15 species (*Astragalus crassicarpus, Argemone albiflora, Asclepias Asperula, Callirhoe leiocarpa, Centaurea americana, Chasmanthium tifolium, Corydalis curvisiliqua, Digitaria californica, Eragrostis trichodes, Herbertia lahue, Linum rigidum, Pavonia lasiopetala, Polytaenia nuttallii, Tradescantia occidentalis, Liatris mucronata*) were placed within under the same genus/family. The center of each plot is the mean value, the other two lines are -/+ standard errors. The colors were coded corresponding to the grouping of phylogenetic positions (Figure 3).

**Figure 2.**
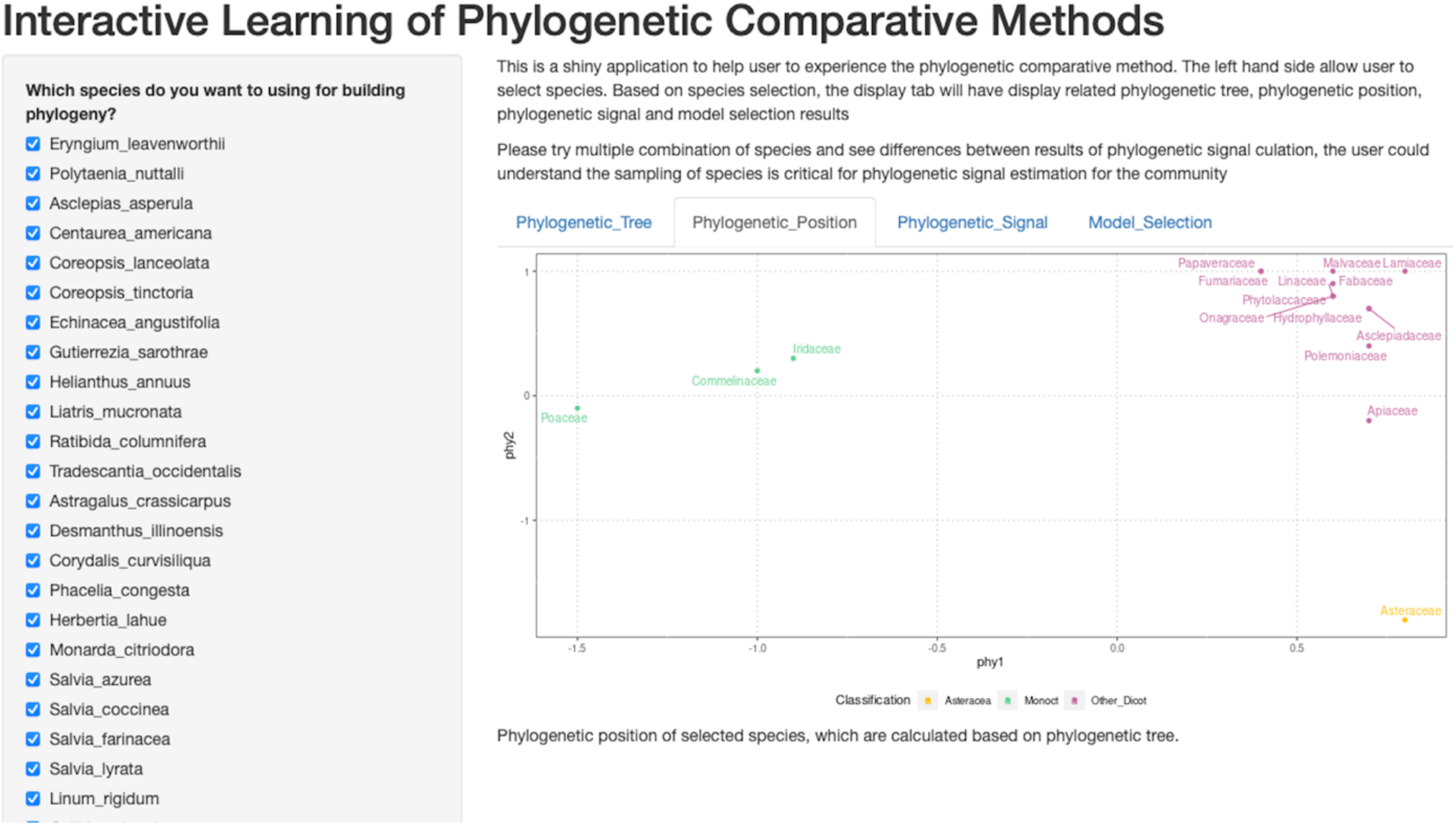
Shiny application of interactive learning of phylogenetic comparative methods. This is a screenshot of the shiny application. The checkbox of species could be used to choose different combinations of species and explore its impact on phylogenetic signals.

### Seed Germination Percentage and Morphological Traits Measurements

Seed germination percentage was obtained from two sources: our own germination trials and previous publications. In all cases, we defined “germination percentage” as the maximum final germination percentage obtained. The germination trials followed a simple germination protocol without cold stratification or other attempts to break dormancy, which simulated minimum requirements for restoration projects. This simple protocol is essentially a measurement of lack of dormancy assuming the tested seeds were full viable. For 34 of the 45 species, we conducted new germination trials. Our new germination trials were trying to simulate the scenario that practitioners want to find some easy to use native species. Because the experiment is trying to simulate the scenario in which practitioners are attempting to find easy to use native species, we bought seeds from a local restoration seed vendor (Native American Seed), and chose species for which they offered local seed sources (and recorded the seed source), with seeds that were harvested less than 6 years ago. When seeds arrived, we stored the seeds in a dry and dark place at room temperature (20 °C) until experiments started. Although it’s possible that some species may exhibit dormancy, we didn’t use any dormancy breaking treatment, in order to simulate simple restoration practice. For the germination experiment, we used triple replicated germination trials: disposable petri dishes with lids were placed in germination chambers (20 °C day and night, with 15 hours and 9 hours day night shift). Inside a petri dish a piece of filter paper was placed to observe auto-claved water to keep the seeds moist. We checked the water sufficiency every day. In each germination trial we split a total of 50 seeds of each species into 5 petri dishes. Since our study used commercial seeds and focused on species dormancy status, we assumed our seeds will either be dormant or start germination within a month. Our observations during experiments proved this assumption. The seeds generally started germinating within 10 days or stayed dormant through the whole germination trial. Our germination trials ran until one week after the last seed germinated. Most of the seed germination trials finished within a month, and all the trials finished within two months. Three independent trials happened in July 2019, September 2019, and November 2019. For the remaining 11 species, we used final germination percentages reported in two published studies (Schwilk and Zavala 2012; Chou *et al*. 2012). These two studies were originally designed for detecting smoke effects on shortgrass prairie species, but we used the control treatment data only which provided conditions similar to those in our trials (20-25 °C, 12-16 hours illumination).

We measured seed mass using an electronic balance (Sartorius Analytical Balance LA 230P, 0.1mg readability) in lab conditions with 10 replicates of 100 seeds each per species. For species in which we could not obtain 100 seeds, we used 30 seeds per replicate.

We measured seed surface area and seed height through digital image processing with 10 replicates. The seed surface was defined by the two largest orthogonal axes, the height was defined as the third axis. We calculated the seed surface area by digital image of the maximum surface area of seeds and imaged under a stereomicroscope at 400 magnification. We transformed the images to 8-bit (black and white) and calculated the surface area using the “analyze particle” function in ImageJ (Schindelin et al. 2012). We also recorded the seeds’ heights calculated by the z-stack image function and NIS-Element BR 4.60.00 software.

### Species Phylogenetic Information

We generated a phylogenetic tree of all study species using two methods: pruning existing phylogeny (Zanne et al. 2014) and binding non-existing tips to the phylogeny based on their taxonomic information. The phylogeny (Zanne et al. 2014) we used in this study was a time-calibrated maximum-likelihood-based phylogenetic tree, built with seven genes (18S rDNA, 26S rDNA, ITS, matK, rbcL, atpB, and trnL-F) downloaded from GenBank. First, we confirmed that every genus in our study was on the Zanne phylogeny. Second, we created a function to prune species which were not on the tree, and we also swapped the species under the same genus if the exact species was not on the tree (see the function of func_prun_replac on https://github.com/chenyanniii/Traits4 repo for more detail). The results showed that 30 species on the tree and 15 missing species (*Argemone albiflora, Asclepias asperula, Astragalus crassicarpus, Callirhoe leiocarpa, Centaurea americana, Chasmanthium latifolium, Corydalis curvisiliqua, Digitaria californica, Eragrostis trichodes, Herbertia lahue, Liatris mucronata, Linum rigidum, Pavonia lasiopetala, Polytaenia nuttallii, Tradescantia occidentalis*). After applying func_prun_replac, 13 of 15 species were placed based in their genus and only two species (*Callirhoe leiocarpa* and *Digitaria californica*) were missing. Thus, we added the missing species (*Callirhoe leiocarpa* and *Digitaria californica*) as sister tips to *Callirhoe involucrate* and *Digitaria ciliaris* under the same genus assuming that phylogenetic relationships were consistent with their taxonomic grouping. Our final tree contained all species was a dichotomous tree (Figure 1).

To incorporate phylogenetic relatedness in the general linear models, we represented the phylogeny by all pairwise phylogenetic distances across taxa. We converted the pairwise distance matrix to points distributed in a two-dimensional coordinate system, using nonmetric multidimensional scaling (NMDS) (`isoMDS` function in the package `MASS`, Venables and Ripley 2002). We evaluated phylogenetic signals for individual traits as Blomberg’s K (Blomberg *et al*. 2003) using the `phylosig` function in the `phytools` R package (Revell 2012). We tested for phylogenetic signal using a randomization test (`phylosig` function) that compared the measured value of Blomberg’s K against a distribution of K calculated when trait values were randomized across the tips of the phylogeny.

### Germinability Prediction Model Selection

To generate and evaluate generalized linear models, we applied backward stepwise model comparison based on the Akaike information criterion (Akaike 1998) using the `AICc` function in the `AICcmodavg` package (Mazerolle 2020). We also used seed germination percentage, three seed morphological traits (seed mass, seed height and seed surface area) and phylogenetic positions to generate a global general linear model. Then, we used AIC to correct for small sample sizes (AICc) and evaluate the fitness of models. We standardized all input parameters to the mean of zero to produce standardized coefficients between parameters for numeric reasons in fitting. We also tested correlation among morphological traits (seed mass, seed height and seed surface area). All original data and scripts that we used to calculate phylogenetic signals, phylogenetic residuals, and seed germinability prediction models are available on GitHub website (https://github.com/chenyanniii/Traits4, DOI: 10.5281/zenodo.6609175).

### Random Sub-sampling of Different Species Pool Size

To estimate the minimum species pool size for obtaining a stable phylogenetic signal, we created 31 different species pool subsets, from 10 species to 40 species. For each pool size, we randomly withdrew 100 times at each pool size species from the whole species pool, thus generating 100 sub-pools of each species pool size by random sub-sampling. The phylogenetic signals of each sub-pool were calculated for their Blomberg’s K and related p value. We analyzed the relationship between sample size and detection rate of phylogenetic signals was analyzed to evaluate the effect of sample size to estimated phylogenetic signals in traits.

### Shiny Application

Shiny is a web framework for displaying data. Shiny is a good data processing demonstration tool, an interactive way for users to experience how different input and procedure affect output. We designed our shiny application to import with our full dataset and display data analysis and results. Users can see our full dataset result (as default), or interactively calculate all parameters for any sub-pools using checkboxes of species (Figure 2).

## Results

In this study, we used 45 commonly selected restoration species to explore the phylogenetic distance among shortgrass prairie species by pruning unnecessary species and adding desired species to the existing phylogenetic tree of flowering plants (Figure 1).

### Seed Final Germination Percentage and Morphological Traits Measurements

When examining species’ trait value with the phylogenetic tree (Figure 1), we found the phylogenetic patterns in seed mass, seed height, seed surface area and seed germination rate were varied. We were not able to germinate eight species (Figure 1, *Argemone albiflora, Callirhoe leiocarpa, Corydalis curvisiliqua, Herbertia lahue, Oenothera rhombipetala, Pavonia lasiopetala, Phytolacca americana*, and *Polytaenia nuttallii*). *Eragrostis trichodes* had the highest final germination percentage, 82%. For seed mass, *Sporobolus airoides* had the lightest weight per seed, 0.0945 ± 0.0083 mg per seed; the heaviest seed was *Pavonia lasiopetala*, 18.75 ± 0.3487 mg per seed. The seed height measurement ranged from 0.658 ± 0.1051 (*Coreopsis tinctoria*) to 2.995 ± 0.1334 mm (*Pavonia lasiopetala*); and the seed surface areas ranged from 0.361 ± 0.0083 (*Sporobolus cryptandrus*) to 25.258 ± 1.322 (*Polytaenia nuttallii*) mm^2^ (Figure 1).

### Species Phylogenetic Information

We used nonmetric multidimensional scaling (NMDS) to compress the phylogenetic distance matrix to a two-dimensional space, with a pressure of 17.86. Our results showed that 45 species were grouped into three clusters: Monocot, Asteraceae and eudicots-except Asteraceae (Figure 3). NMDS compressed phy1 (x-axis) corresponded to separating monocot and eudicots, while the phy2 (y-axis) separated Asteraceae from other families.

**Figure 3.**
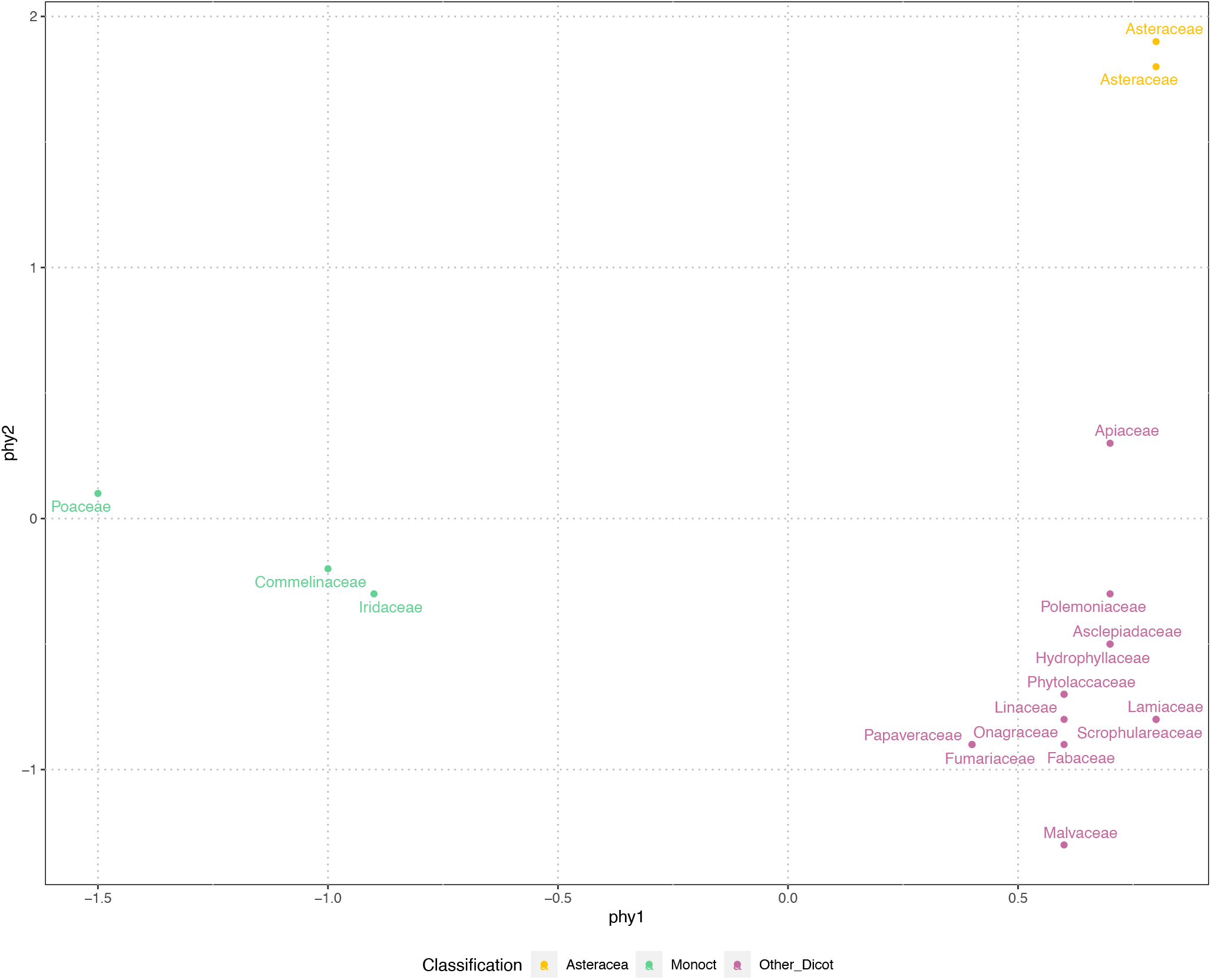
Phylogenetic position of 45 species, represented by family, were clustered in three groups. The phylogenetic positions were generated from paired-wise distances of species on the phylogenetic tree (see Figure 1). The nonmetric multidimensional scale (NMDS) was applied, at the stress of 17.86, displayed in two axes. For the convenience of display the phylogenetic positions were grouped and color coded by vision.

Our measurements of phylogenetic signals, Blomberg’s K (using species shuffling method), were low for all four seed traits, indicating a departure from signal under strict Brownian motion and suggesting that these traits are evolutionarily labile. Although Blomberg’s K were low, indicating a weak phylogenetic signal, we found significant phylogenetic signals for seed mass (K = 0.07, p = 0.01) and seed height (K = 0.05, p = 0.05) (Table 2).

**Table 2.** Phylogenetic signal was tested in seed morphological traits and overall seed final germination percentage. Blomberg’s K was used to evaluate phylogenetic signals (Blomberg et al. 2003).

| Trait | Blomberg's K | P-value |
| --- | --- | --- |
| Seed Mass | 0.07 | 0.01 |
| Seed Height | 0.05 | 0.05 |
| Seed Surface Area | 0.03 | 0.14 |
| Seed Final Germination Percentage | 0.02 | 0.20 |

K = 1, the traits is perfectly fit with Brownian motion model
K > 1, the traits is more conserved than expected comparing to Brownian motion model
K < 1, the traits is less conserved than expected comparing to Brownian motion model
\* indicate traits containing phylogenetic signal (P =< 0.05)

### Germinability Prediction Model Selection

The full set of models built from morphological traits and phylogenetic information were evaluated using adjusted AIC (AICc). The AICc values range from 129.9 to 139.4. The best prediction model is using seed height to predict seed germination (AICc = 129.9), slightly better than the model using seed mass to predict germination (AICc = 130.5). The models with low AICc values were clustered by using one morphological trait as a predictor or the combination of two morphological traits. This indicated that morphological traits out-perform phylogenetic distance in predicting seed germination. Pearson correlation coefficient analysis revealed a strong correlation between seed mass and seed height (r = 0.66, p < 0.01); a medium correlation between seed mass and seed surface area (r = 0.49, p <0.01); no correlation was detected between seed height and seed surface area.

### Random Sub-sampling of Different Species Pool Size

We calculated phylogenetic signals of morphological traits (seed mass, seed height, and seed surface area) and seed germination rate of all 3,100 sub-pools. All Blomberg’s K values were between 0 and 1 in all phylogenetic signal calculations, except 9 of them were larger than 1. In general, phylogenetic signals distributed widely at small species pool sizes, and became less varied while increasing species pool sizes (Figure 4). For seed height, seed surface area, and seed germination, the probability of detecting phylogenetic signals were consistently low regardless of the species pool size. This was true even for seed height, for which we detected a significant phylogenetic signal in our full dataset. In contrast, the probability of detecting the phylogenetic signal of seed mass increased with species pool size (Figure 5).

**Figure 4.**
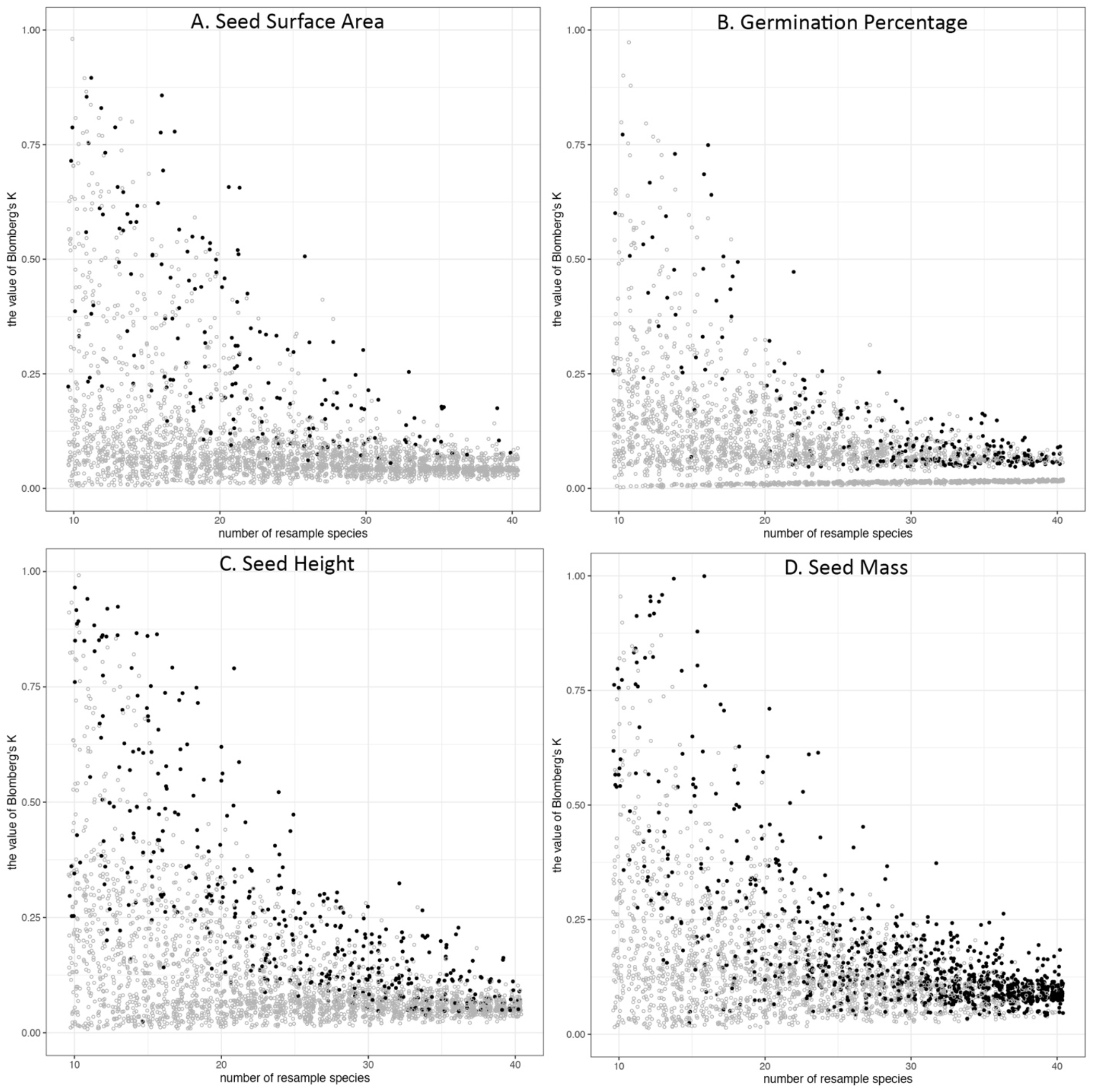
The distribution of Blomberg’s K along the size of the species pool in random subsampling tests. The species were resampled 100 times from 10 species to 40 species, and phylogenetic signal (Blomberg’s K) was calculated for each trait, 3100 times for each trait. Phylogenetic signals of (A) seed surface area, (B) germination percentage, (C) seed height, (D) seed mass. The dots represent the Blomberg’s K value of each resampling pool. The color of dots indicates the p-value of Blomberg’s K (p < = 0.05, black; p > 0.05, grey).

**Figure 5.**
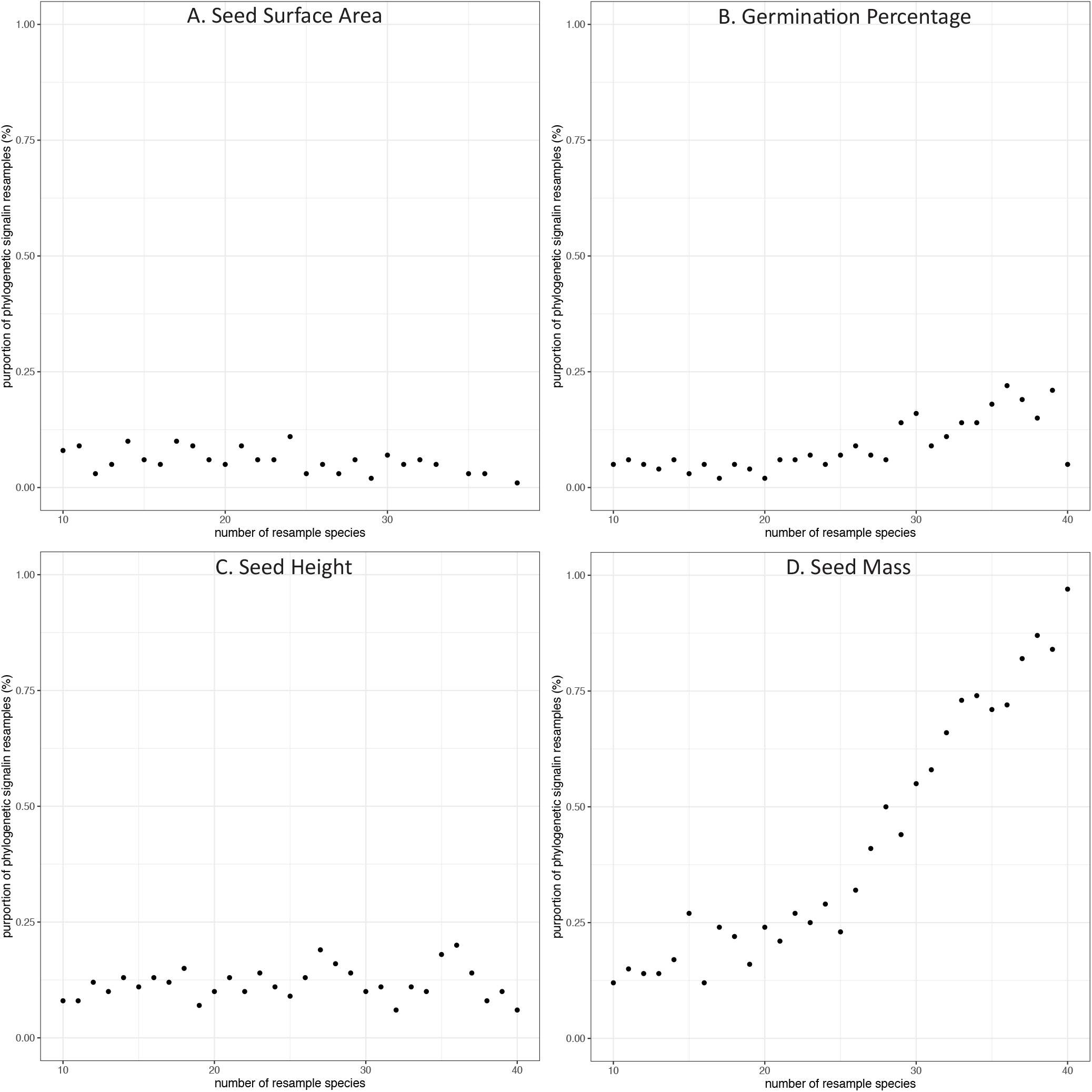
The proportion of subsamples with significant phylogenetic signals along the change of number of species in species pools. The species were resampled 100 times from 10 species to 40 species. The dots represent the proportion of Blomberg’s K value (p =< 0.05) in each resampling pool: (A) seed surface area, (B) germination percentage, (C) seed height, (D) seed mass.

## Discussion

Aiming to verify the usefulness of trait conservatism in restoration seed selection, we measured seed traits, ran seed germination tests, calculated phylogenetic signals in seed traits, and presented the phylogenetic residual in seed germinability prediction models. We quantified weak phylogenetic signals in seed mass and seed height, but we found no phylogenetic signal in seed surface area nor in seed final germination percentage. In those traits that did exhibit phylogenetic signals, the signals were weak: closely related species were more similar than expected under species shuffling, but more different in their trait values than expected under Brownian motion.

### Phylogenetic Tree

The phylogenetic tree of 45 commonly selected species in shortgrass prairie ecological restoration was clustered in Poaceae within monocots and were relatively clustered in Asteraceae and Lamiaceae within eudicots (Figure 3), which reflects that the species composition may be clustered in shortgrass prairie. The phylogenetic comparative methods displayed trait values indicated that the closely related species had similar trait values in seed mass and seed height, but not in seed germination (Figure 1). The NMDS compressing phylogenetic distance into two-dimensions shows three distinct clusters (Figure 3). The results showed that shortgrass prairie families were grouped into 3 clusters: one monocot group and two eudicot groups (Asteraceae and others, Figure 3). Meanwhile the tallgrass prairie species (Barak *et al*. 2018) were grouped into 4 clusters: one monocot group, three eudicot groups (Asteraceae, Fabaceae, and others).

Our development of the Shiny application demonstrated: (1) the procedure of pruning the synthetic phylogenetic tree (Zanne et al. 2014) to the desired species tree (Figure 1). (2) the calculation of compressing phylogenetic distance into two-dimensions. The interactive demonstration allows users to select all or a portion of desired species and understand the effect of species selection on phylogenetic calculation.

### Phylogenetic Signal in Traits

Phylogenetic signal indicates that closely related species have more similar trait value than expected under species shuffling across tips of a phylogeny. We found significant phylogenetic signals in seed mass and seed height, but no such signals in seed surface area nor in seed final germination percentage. Although germination traits are not specific or constant in each species (but vary in space and time), since we chose seeds from the same eco-region, our results are able to represent our region and still allow generalization when considering germinability predictions. Generally, seed mass is phylogenetically conserved in sample taxas from different ecosystems (tallgrass prairie, Barak et al. 2018; alpine grassland, Bu *et al*. 2016; and globally, Westoby et al. 2002). In our set of taxa, we found a weak but significant pattern. Seed mass often predicts energy and nutrient provisioning (Westoby *et al*. 2002), which increases seed germination rates and stress tolerance (Leishman 2000; Moles 2018). This assumes, however, that mass is primarily the embryo and nutrients. It is possible for a large portion of the seed mass to be seed defense structures (i.e. seed coat).

We used seed height and seed surface area as proxies for seed dispersal syndrome, because these dimensions influence primary wind dispersal (seed departure from mother plants, Sheldon and Burrows 1973; Jongejans and Telenius 2001) and secondary wind dispersal (seed lifting off ground by wind power, van Tooren 1988; Schurr *et al*. 2005; Zhu *et al*. 2022). Primary dispersal is mainly related to dispersal height and terminal falling velocity, which is influenced by seed morphology (Sheldon and Burrows 1973; Jongejans and Telenius 2001). Secondary dispersal distance strongly depends on the lift-off velocity, which is influenced by seed height and the planform area of a seed exposed to airflow (van Tooren 1988; Schurr *et al*. 2005; Zhu *et al*. 2022). Classically, seed shape was measured by the roundness or closeness of a seed to specific shape, such as ellipse or cardioid (Cervantes et al. 2016) and linked with seed persistence in soil seed bank (Moles *et al*. 2000; Laughlin 2014). Some recent studies link seed morphological shape with evolutionary constraint and selective pressure of seeds and its potential relationship with seed germination (Barak et al. 2018, Bu *et al*. 2016). In our study, seed mass and seed height were positively correlated. We found a weak pattern of phylogenetic trait conservatism in two traits, but this signal did not aid in improving seed germinability prediction models.

Seed germination is a complex phenomenon. Our measure of total germination was, in effect, a dormancy proxy: high germination rates indicated a lack of dormancy in our research. Our experiment didn’t include any dormancy breaking retreatments, only supplying light and water during experiments to simulate practitioners’ low effort practices. Seed germination can be influenced by abiotic factors, such as wetland species germination impacted by water level (Keddy 1992); or arid zone woody species developing rapid germination in response to unpredictable rainfall (Duncan *et al*. 2019). Seed germination can also be influenced by biotic factors, such as small- and large-seeded species diverging in the species they associate with, regarding seed mass and understory light preference (Umaña *et al*. 2020). We didn’t detect a phylogenetic signal in germination rate indicating this trait is highly labile. This result was different from a similar study of tallgrass prairie species (Barak *et al*. 2018), where the authors found significant phylogenetic trait conservatism in germination percentage under control and gibberellic acid treatment, and including phylogeny improve time-to-germination (survival) model. However, the survival model (Barak *et al*. 2018) includes both germination time and pretreatment for germination rate and doesn’t measure dormancy. Differing patterns in phylogenetic signal in germination rate of two prairie studies are reasonable, in consideration of environmental differences between two different ecosystems, and the germination experiment setting in two studies.

### Germinability Prediction Model Selection

The germinability predictive models with morphological data did not improve when adding phylogenetic information using the full dataset (Supplemental Material). This means adding phylogenetic information to morphological measurements increased the complexity of models but did not increase the fitness of models. This is not surprising given that we found no phylogenetic signal in seed germination rate and only weak signals in two other traits.

### Random Sub-sampling of Different Species Pool Size

From the distribution of Blomberg’s K, we can tell the species sample size will greatly influence phylogenetic signal calculation (Blomberg *et al*. 2003). Our shortgrass prairie restoration species results showed that the phylogenetic signal would be less impacted by the species composition, and less varied with sufficient species, around 35 to 40 (Figure 4). This also indicates the 45 species we have in our study is sufficient.

In the full dataset (45 species), we were able to detect phylogenetic signals for both seed mass and seed height. However, the subsampling exploration method demonstrates that detecting a phylogenetic signal in seed height is a low probability event. On the other hand, our sub-sampling in seed mass showed that the probability of detecting a phylogenetic signal increased along with the increase in the number of species in the species pool. The Blomberg’s K value is stable at 40 species, which could indicate that if researchers or practitioners have over 40 species sub-sampling of shortgrass prairie restoration species, their studies should be able to detect phylogenetic signals. The random sampling methods to verify sample size method could apply in sampling species to estimate phylogenetic conservatism in plant communities.

### Shiny Application

From the Shiny application, restoration practitioners could use interactive methods to explore our data and statistical analysis and results visualization. For readers who are first exposed to phylogenetic comparative methods, the interactive graphic user interface can lower the bar for exploring our data, as well as increase engagement. Our checkbox of species list allows users to design their composition of species, and to investigate the impact of species choice on phylogenetic signal and germinability prediction. Our Shiny application was published on GitHub website (https://github.com/chenyanniii/Traits_Shiny, DOI: 10.5281/zenodo.6609191) and on shinyapps.io (https://chenyanniii.shinyapps.io/Phylo_Compar_Traits/).

### Comparison Between Tallgrass Prairie and Shortgrass Prairie Studies

Seed germination is a complex physiological phenomenon that could be studied for its optimization using dormancy breaking treatments (Barak et al. 2018), as well as could be a dormancy proxy, such as high germination rates indicated a lack of dormancy in our research. Our research can be contrasted with a was similar tallgrass prairie study (Barak et al. 2018), in which: (1) the phylogenetic signals of germination were detected in morphological traits and seed germination percentage; (2) phylogenetic information improves the seed germinability prediction model. We saw the potential of applying phylogenetic information in ecological restoration, so we tested the phylogenetic application in simply restoration setting: (1) We selected regional appropriated seed sources from a local restoration vendor. (2) We proxy dormancy in seed sources by running germination trails without any dormancy breaking treatment to approximate the conditions preferred by restoration practitioners. (3) We tested our results against null models: confirming our confidence in sample size, examining the robustness of our conclusion while ensuring we can generalize results for the whole shortgrass prairie plant community. Our unique restoration scenario of shortgrass prairie showed a few advancements of knowledge. First, only seed mass and seed height detected phylogenetic signals in 45 species.

The phylogenetic signal in seed mass is well preserved and can be generalized to estimate the phylogenetic signal for the shortgrass prairie plant community. On the opposite, detecting a phylogenetic signal in seed height is a low chance event that the phylogenetic signal in 45 species should not be generalized to estimate the phylogenetic signal for the shortgrass prairie plant community. Second, estimating phylogenetic signals for a plant community needs a larger sample size than a single fixed group. The shortgrass prairie plant community needs at around 40 species for detecting a general pattern (Figure 4 and Figure 5), which is twice of the twenty species assumption in a fixed species comparative study (Blomberg et al. 2003).

## Conclusion and Future Studies

Overall, we have demonstrated that the phylogenetic signal calculation can be influenced by size and composition of seed pool. We recommend running a sub-sampling test to verify the sufficiency of species and phylogenetic conservatism in traits for a community study, and we proposed a general protocol for implementing phylogenetic conservatism in plant community restoration (Figure 6). Our Shiny application is on GitHub website (https://github.com/chenyanniii/Traits_Shiny, DOI: 10.5281/zenodo.6609191) and on shinyapps.io (https://chenyanniii.shinyapps.io/Phylo_Compar_Traits/), using an interactive way to demonstrate how species composition directly impacts the phylogenetic signal calculation.

**Figure 6.**
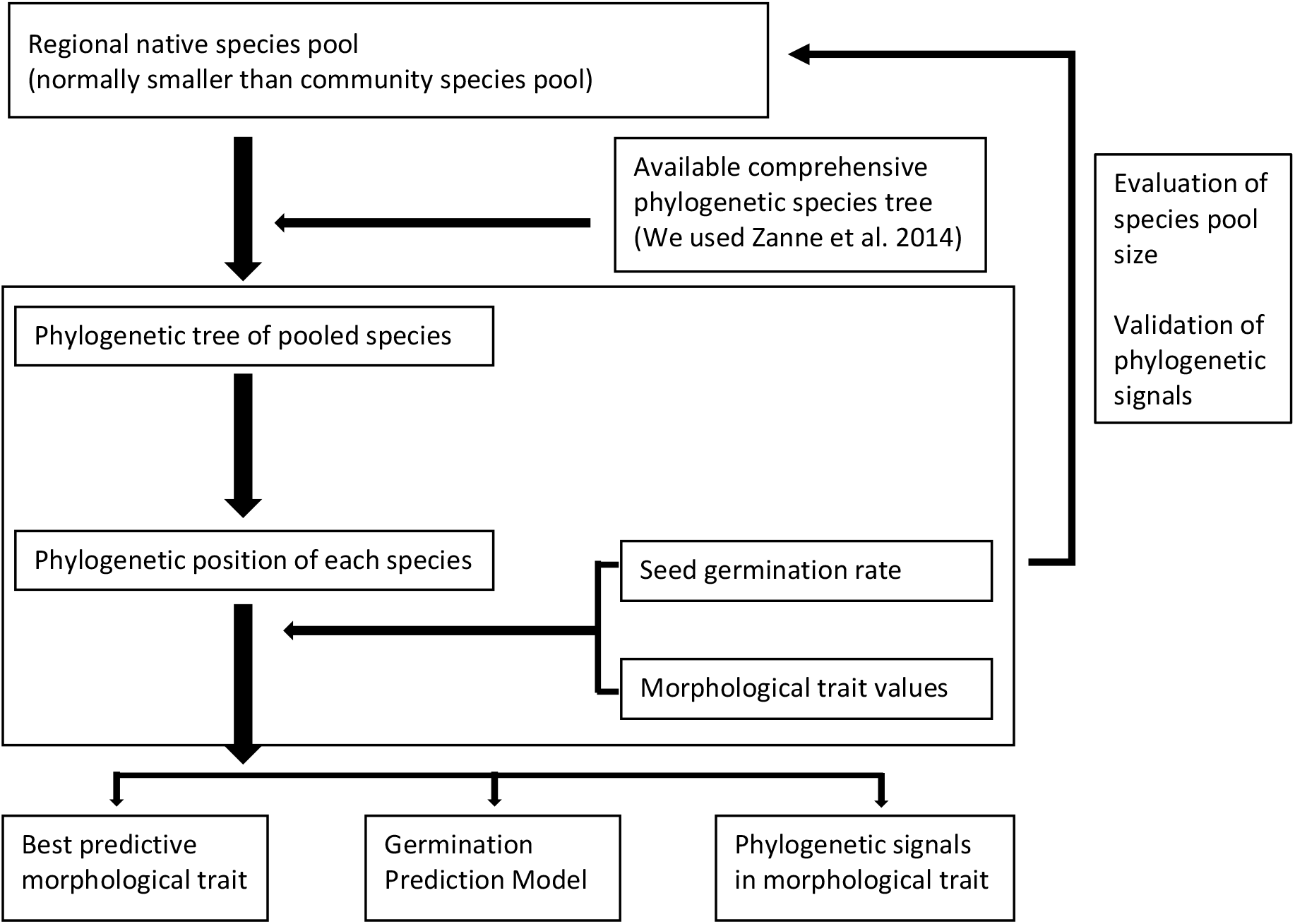
General protocol for generating a germinability prediction model with phylogenetic information for a plant community. This model needs a pool species with phylogenetic information, morphological data and germination data to build. It will be able to explore the germination pattern of the community.

Our work demonstrated that morphological traits (seed mass and seed height) are highly conserved traits in shortgrass prairie, North America. Yet our study could not detect the benefit of adding phylogenetic information using morphological traits to predict seed germination. The inconsistent role of phylogeny in different ecosystems needs further exploration, especially taking advantage of large standard databases of seed traits and the tree of life.

## Glossary

### Phylogeny / Phylogenetic tree

branching evolutionary histories / to graphs that represent these evolutionary histories. Phylogenetic tree including gene tree and species tree. In this paper, we only refer to species’ tree (Baum and Smith 2013).

### Phylogenetic conservatism

the hypothesis that closely related species share more traits than distantly related species. (Agrawal 2007).

### Phylogenetic position

the relative position between species commonly used nearest neighbor and paired-wise distance. We used paired-wise distance in our calculation.

### Phylogenetic signal

to describe a tendency for evolutionarily related organisms, under assumption of following a certain evolutionary model, to resemble each other. (Blomberg *et al*. 2003).

### Phylogenetic residual

incorporate the phylogeny through error structure, such as estimating ancestral states, rates of evolution, phylogenetic effects. (Garamszegi 2014).

### Supplemental Material

Model Evaluation for all seed germinability prediction models. AICc is the adjusted AIC value due to the small sample size in biological tests.

