## Supplemental Table 1 for "Including Phylogenetic Conservatism of Shortgrass Prairie Restoration Species Does Not Improve Species Germinability Prediction"

| | Model | AICc |
|---|---|---|
| 1 | scaled_Germination ~ scaled_Height | 129.89 |
| 2 | scaled_Germination ~ scaled_Mass | 130.52 |
| 3 | scaled_Germination ~ scaled_Area + scaled_Height | 131.68 |
| 4 | scaled_Germination ~ scaled_Height + scaled_Mass | 131.93 |
| 5 | scaled_Germination ~ scaled_Height + phy1 | 132.2 |
| 6 | scaled_Germination ~ scaled_Height + phy2 | 132.23 |
| 7 | scaled_Germination ~ scaled_Area | 132.42 |
| 8 | scaled_Germination ~ scaled_Mass + phy2 | 132.62 |
| 9 | scaled_Germination ~ phy2 | 132.8 |
| 10 | scaled_Germination ~ scaled_Area + scaled_Mass | 132.92 |
| 11 | scaled_Germination ~ scaled_Mass + phy1 | 132.93 |
| 12 | scaled_Germination ~ phy1 | 133.18 |
| 13 | scaled_Germination ~ scaled_Area + scaled_Height + phy2 | 134.05 |
| 14 | scaled_Germination ~ scaled_Area + scaled_Height + phy1 | 134.08 |
| 15 | scaled_Germination ~ scaled_Area + phy2 | 134.14 |
| 16 | scaled_Germination ~ scaled_Area + scaled_Height + scaled_Mass | 134.18 |
| 17 | scaled_Germination ~ scaled_Height + scaled_Mass + phy1 | 134.35 |
| 18 | scaled_Germination ~ scaled_Height + scaled_Mass + phy2 | 134.36 |
| 19 | scaled_Germination ~ scaled_Height + phy1 + phy2 | 134.68 |
| 20 | scaled_Germination ~ scaled_Area + phy1 | 134.79 |
| 21 | scaled_Germination ~ scaled_Area + scaled_Mass + phy2 | 135.09 |
| 22 | scaled_Germination ~ phy1 + phy2 | 135.11 |
| 23 | scaled_Germination ~ scaled_Mass + phy1 + phy2 | 135.15 |
| 24 | scaled_Germination ~ scaled_Area + scaled_Mass + phy1 | 135.45 |
| 25 | scaled_Germination ~ scaled_Area + scaled_Height + phy1 + phy2 | 136.61 |
| 26 | scaled_Germination ~ scaled_Area + phy1 + phy2 | 136.63 |
| 27 | scaled_Germination ~ scaled_Area + scaled_Height + scaled_Mass + phy2 | 136.69 |
| 28 | scaled_Germination ~ scaled_Area + scaled_Height + scaled_Mass + phy1 | 136.71 |
| 29 | scaled_Germination ~ scaled_Height + scaled_Mass + phy1 + phy2 | 136.93 |
| 30 | scaled_Germination ~ scaled_Area + scaled_Mass + phy1 + phy2 | 137.75 |
| 31 | scaled_Germination ~ scaled_Area + scaled_Height + scaled_Mass + phy1 + phy2 | 139.38 |
